# Duplex Reverse Transcription Loop-Mediated Isothermal Amplification on a Nanofluidic Digital Chip (Nano-dChip)

**DOI:** 10.64898/2026.03.18.712394

**Authors:** Natalie Luu, Li Liu, Esmeralda Ruiz-Garcia, Juhong Chen, Steve J. Dollery, Gregory J. Tobin, Ke Du

## Abstract

Over the past decades, the frequency of viral outbreaks has increased substantially worldwide, driven in part by global travel and resulting in millions of deaths each year. This trend underscores the urgent need for rapid, simple, and accessible diagnostic tools for infectious disease detection. Here, we present a nanofluidic digital chip (Nano-dChip) for point-of-care viral RNA detection that delivers results within 30 minutes at a cost of less than $0.50 per chip. The Nano-dChip employs reverse transcription loop-mediated isothermal amplification (RT-LAMP) for highly sensitive and specific target amplification. Reaction reagents are compartmentalized into numerous nanofluidic reservoirs, enabling highly quantitative detection while minimizing contamination risks. Using a single chip, we successfully detect both SARS-CoV-2 and Influenza H3 RNA with a detection limit of 10 fM, demonstrating the Nano-dChip’s potential as a rapid, low-cost, and scalable diagnostic platform for timely outbreak control.

## A. Introduction

In recent years, isothermal amplification methods such as recombinase polymerase amplification (RPA) and loop-mediated isothermal amplification (LAMP) have gained significant attention as alternative diagnostic approaches to polymerase chain reaction (PCR) [1,2]. These methods enable nucleic acid amplification at a constant temperature, eliminating the need for a thermocycler and thereby enhancing their suitability for point-of-care (POC) applications. LAMP employs four to six primers that recognize multiple regions of a target gene, providing high specificity and robust amplification [3]. An extension of this method, reverse transcription LAMP (RT-LAMP), enables the direct amplification of RNA targets by incorporating reverse transcriptase into the reaction, eliminating the need for a separate reverse transcription step. RT-LAMP has demonstrated sensitivity comparable to that of RT-PCR and has been widely used for detecting SARS-CoV-2 and other viral pathogens [4,5], highlighting its potential as a rapid, reliable, and field-deployable diagnostic tool.

We recently developed a nanofluidic digital chip (Nano-dChip) that integrates RT-LAMP for the rapid detection and quantification of viral RNA [6,7]. The device is fabricated by bonding a PDMS chip containing more than 1,000 individual reservoirs onto a glass slide, creating a simple and low-cost platform. By miniaturizing reactions within a nanofluidic format, the Nano-dChip reduces reagent consumption, minimizes cross-contamination, and enables highly parallelized analysis. At low concentrations, the users only need to count number of the fluorescent reservoirs, enabling simple and highly quantitative nucleic acid detection.

In this study, we further demonstrate duplex, quantitative detection of two respiratory viruses— SARS-CoV-2 and Influenza H3 RNA—on a single Nano-dChip by introducing two sets of virus-specific primers into the device. As shown in **Figure 1a**, the SARS-CoV-2 primer was labeled with a 5′ 6-FAM dye (IDT) while the Influenza H3 primer was labeled with a 5′ Cy5™ dye (IDT).

**Figure 1.**
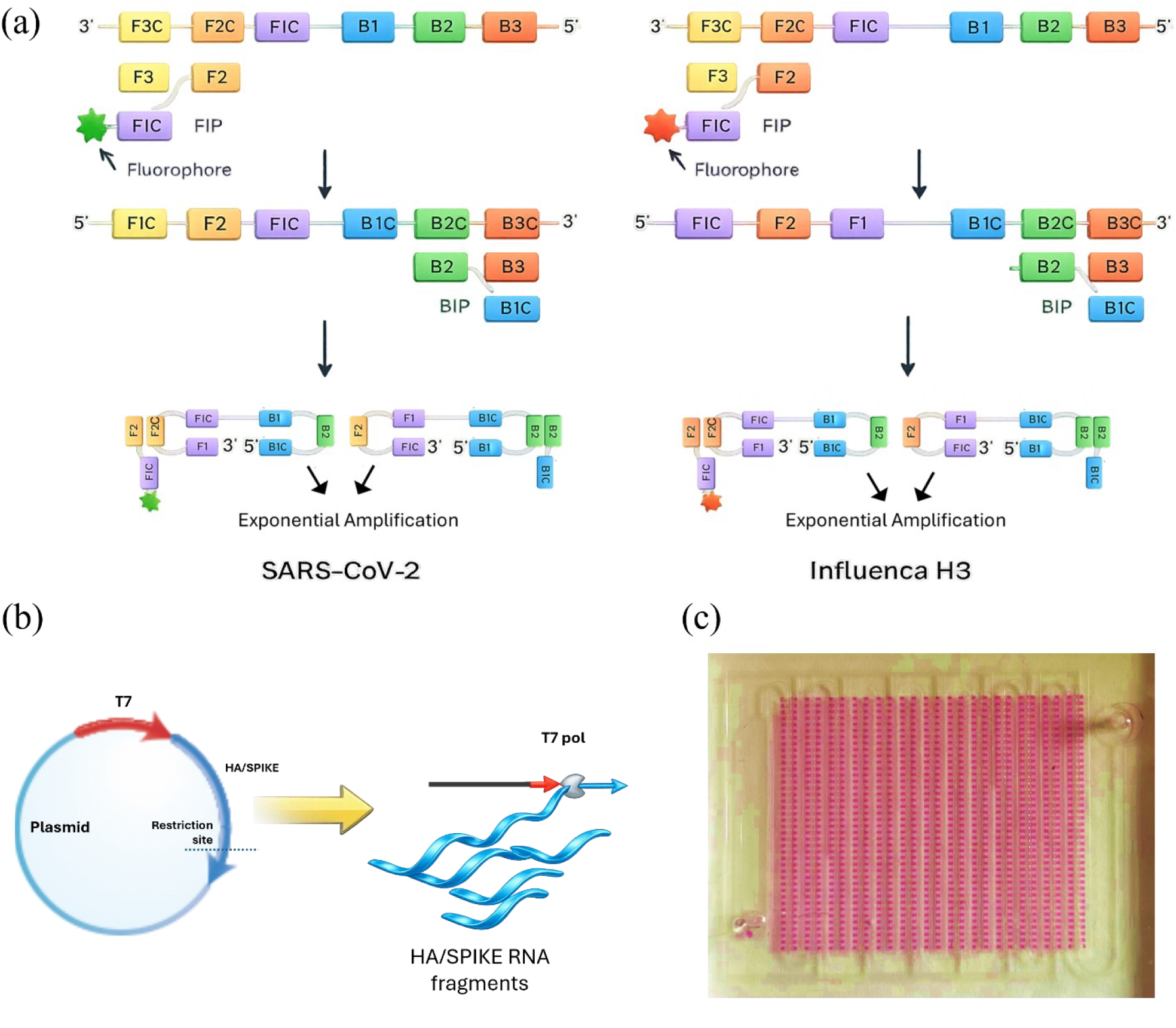
Schematic of the duplex RT-LAMP reaction for the simultaneous detection of both SARS-CoV-2 (green channel) and Influenza H3 (red channel). The sequence specific exponential amplification incorporates the fluorophore in the amplicon. (b) Schematic for the preparation of SARS-CoV-2 and Influenza H3 target. (c) Top view of the Nano-dChip filled with phenol red dye and each well separated by mineral oil.

Duplex detection was achieved by switching between the green and red fluorescence channels. The results were further confirmed by bright-field imaging: in the presence of both targets, the reaction mixture turned yellow, enabling a simple visual readout without the need for imaging instrumentation. Importantly, we achieved a detection limit of 1 fM for both targets with a dynamic range spanning five orders of magnitude, underscoring the potential of this platform for developing a simple, portable POC diagnostic system [8].

## B. Experiments

### Preparation of SARS-CoV-2 and Influenza H3 RNA targets

To linearize plasmids for transcription, 6 µg of P1041 HA-WT-pTriEx was digested with SpeI, and 6 µg of P2442 Cov2 Spike plasmid was digested with HindIII (New England Biolabs, MA, USA) [9]. Linearization was confirmed by PAGE analysis of 0.25 µg DNA. The remaining DNA was ethanol precipitated twice and resuspended in RNase-free water. RNA was synthesized using the T7 RiboMAX™ RNA production kit (Promega, WI, USA) in 100 µL reactions for 4 h at 37 °C, followed by 30 min DNase treatment to remove template DNA according to the manufacturer’s instructions (RiboMAX™, Promega, WI, USA). Resulting RNA was purified using silica membrane column purification (RNeasy™, QIAGEN, MD, USA). RNA products were analyzed by PAGE using a RiboRuler RNA ladder (Thermo Fisher Scientific, MD, USA) and quantified by UV absorbance at 260 nm. The schematic of the target preparation is shown in **Figure 1b**.

### Fabrication of Nano-dChip

The fabrication of the Nano-dChip has been reported by us before. Briefly, SU-8 2075 negative photoresist was used to create the mold for the PDMS chip. The photoresist was spin coated on a new silicon wafer, followed by soft baked on a hot plate at 65°C for 5 minutes and at 95°C for10 minutes. After UV exposure, post exposure bake was performed at 65°C for 5 minutes then at 95°C for 10 minutes. The mold was developed in SU-8 developer for 10 minutes, rinsed with isopropyl alcohol for 10 seconds, and dried with pressurized air. Before PDMS molding, salinization was performed for 30 minutes and PDMS base and curing agent with a ratio of 10:1 was poured on the mold, degassed once more in the vacuum before being placed into an oven at 60°C for 2 hours. After peeling off the PDMS slab from the mold, holes were punched and it was boneded to a glass slide with oxygen plasma (Electro-Technic Products) for 1 minute. The micrograph of a Nano-dChip filled with red dye is shown in **Figure 1c**.

### Duplex on-chip experiments

A mixture consisting of 25 μL of WarmStart® Colorimetric LAMP 2X Master Mix (New England Biolabs), 5 μL of LAMP primer (Integrated DNA Technologies) mix at a 10X concentration, 1.2 μL of quencher probe (Integrated DNA Technologies) at 100μM, 14.8 μL of distilled water (dH2O) and 4 μL of the target RNA were combined in a 600 μL microcentrifuge tube. Half of the volume of the primer mix, quencher probe, and target RNA were those designed for virus 1, whereas the other half of the volume was those designed for virus 2. Five microliter of phenol red dye was added to the mixture, resulting in a total volume of 55 μL. The phenol red dye was used to enhance the contrast of the fluid to the background and track the flow path of the mixture as it is fed into the chip. Before sample loading, the Nano-dChip was vacuumed for 15 minutes, allowing the air inside the reservoirs and main channels to be diffused into the vacuum lungs through the permeable PDMS membrane. After loading the mixture into the reservoirs, mineral oil was flowed into the channels to prevent the mixture inside the reservoirs from evaporating during incubation. The chip was placed on a hot plate at 64 °C for 30 minutes to initiate the LAMP reaction.

## C. Results and Discussion

To evaluate the feasibility of the duplex LAMP assay, we compared no-template controls, negative controls, and positive controls using a target concentration of 1 nM (**Figure 2**). The top row shows the no-template control, which contains the complete LAMP reagent mixture but lacks target RNA. Under bright-field imaging, only droplets are observed in each well, with no visible color change. Consistently, images acquired using the FAM and Cy5 filters show no detectable fluorescence, confirming a low background signal. In the negative control experiments, SARS-CoV-2 targets were added to the Influenza H3 LAMP reagents, and Influenza H3 targets were added to the SARS-CoV-2 LAMP reagents. No fluorescence signal was detected in either channel, demonstrating the high specificity of the primer design. In contrast, for the positive controls, the corresponding target sequences were added to their respective LAMP reagents. The reagents under bright field image turn to yellow color after the reaction. In addition, both green (FAM) and red (Cy5) fluorescence signals were simultaneously detected on the same chip, confirming successful duplex detection. After validating the duplex LAMP assay, we evaluated the detection sensitivity for both the SARS-CoV-2 and Influenza H3 targets over a concentration range from 1 fM to 100 pM (**Figure 3**). No color change was observed at 1 fM. At 10 fM, distinct green and red reservoirs appeared, indicating successful detection of both targets at low concentrations. As the concentration increased to 100 fM, a greater number of colored reservoirs was observed, consistent with increased amplicon generation during the LAMP reactions. Starting at 1 pM, yellow reservoirs began to appear, indicating the co-localization of both targets within the same reservoirs. The proportion and uniformity of yellow reservoirs further increased at higher concentrations, suggesting that most reservoirs contained both targets. Fluorescence imaging revealed a concentration-dependent increase in signal intensity for both targets. In the FAM channel, the number of green fluorescent reservoirs increased progressively from 10 fM to 100 pM, a trend that was consistent with the Cy5 channel. These results demonstrate that the platform enables quantitative duplex detection across a wide dynamic range.

**Figure 2.**
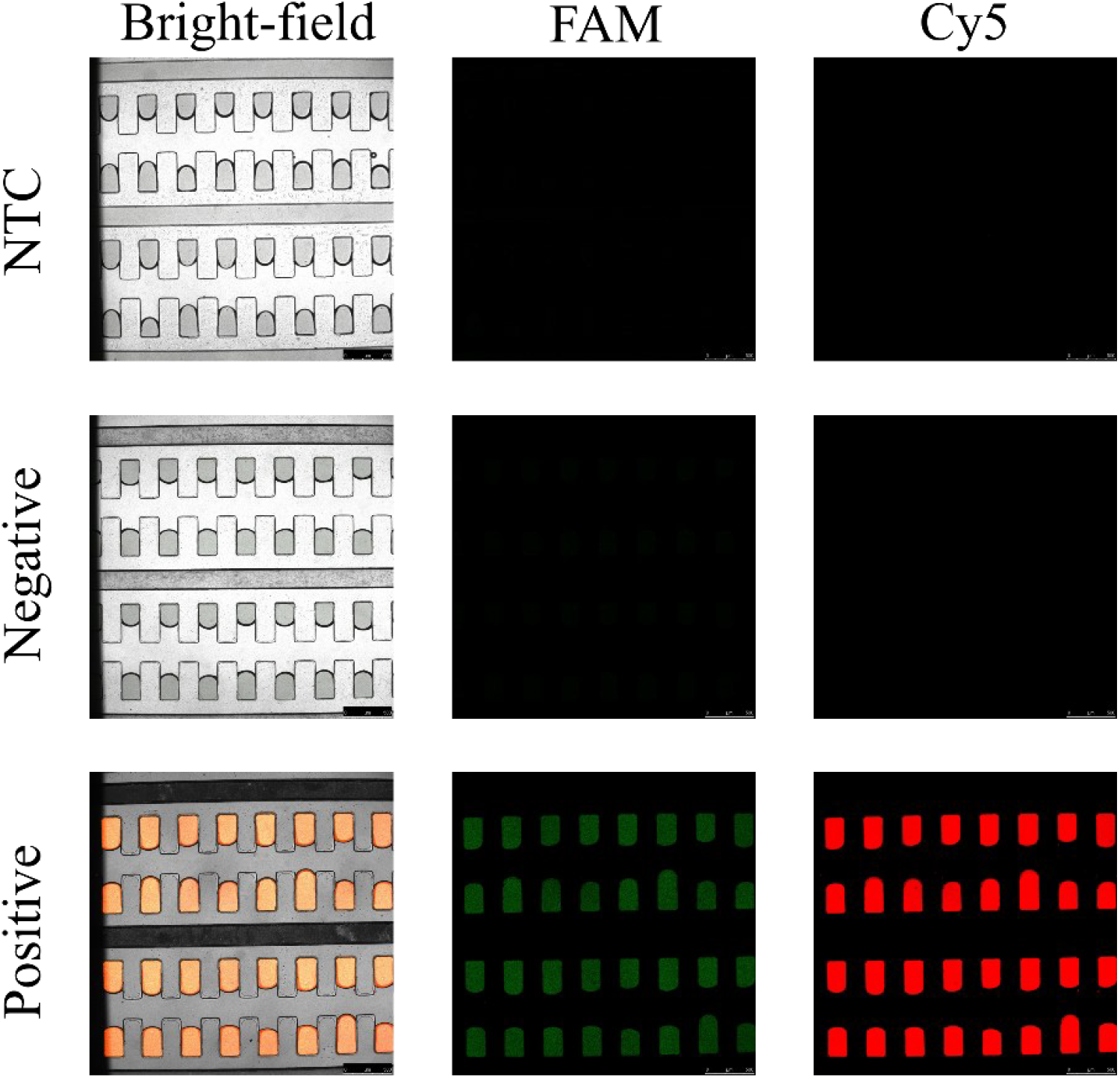
Confocal microscopy images (bright-field and fluorescence) of Nano-dChip micro reservoirs following duplex RT-LAMP detection of SARS-CoV-2 and Influenza H3 RNA. NTC (no template control): Complete RT-LAMP reaction mixture without target RNA. Negative control: RT-LAMP reagents combined with a non-matching (incorrect) target. Positive control: RT-LAMP reagents with the corresponding target RNA at a concentration of 1 nM.

**Figure 3.**
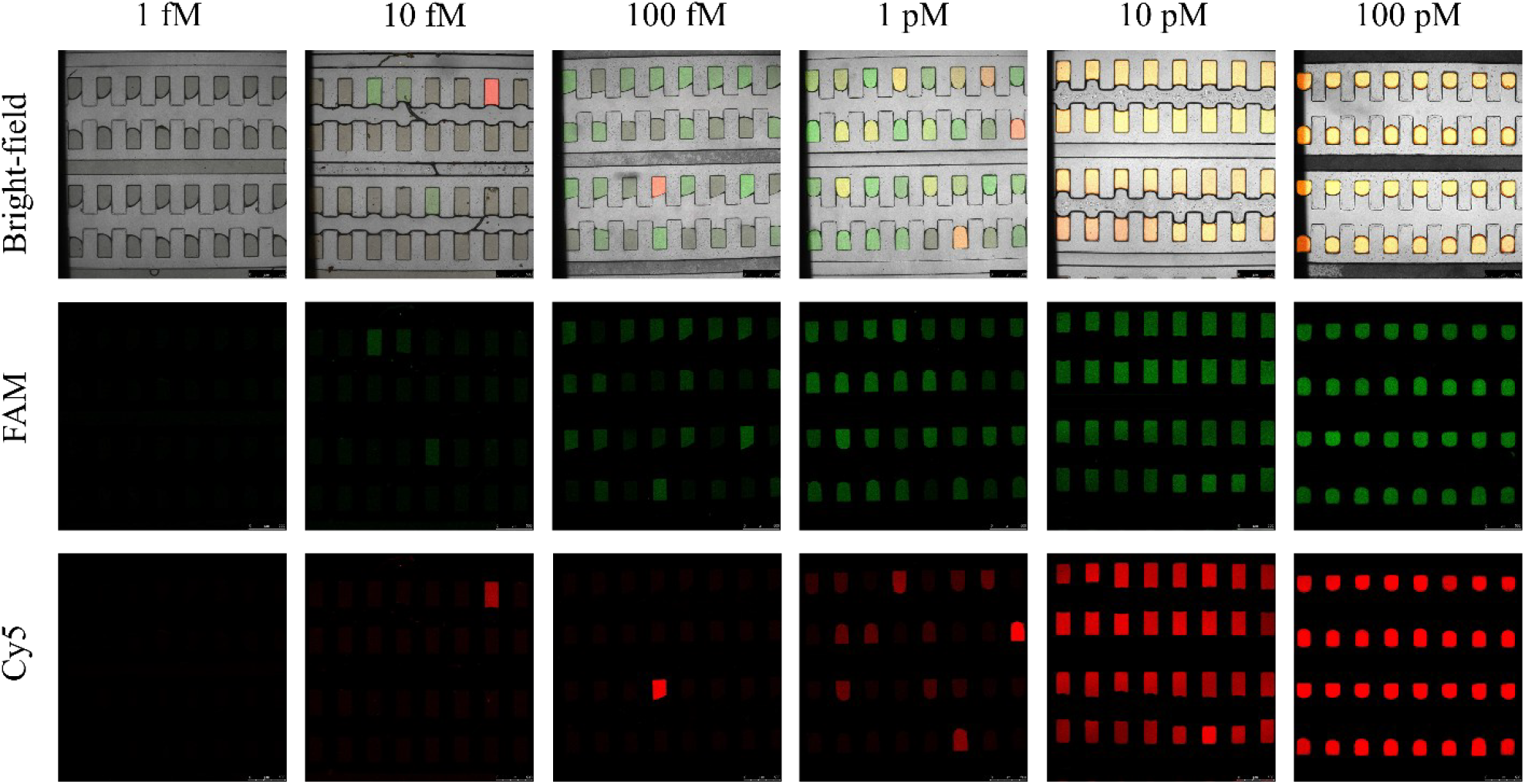
Confocal microscopy images of Nano-dChip reservoirs after RT-LAMP test of both SARS-CoV-2 and Influenza H3 with a target concentration ranging from 1 fM to 100 pM. The top row shows bright-field images of the reservoirs, the middle row shows the green fluorescence channel, and the bottom row shows the red fluorescence channel.

Figure 4 quantifies the fluorescence response of the duplex LAMP assay across a wide concentration range for both SARS-CoV-2 and Influenza H3 targets. As shown, the no-template control (NTC) and 1 fM samples exhibit minimal fluorescence, confirming a low background signal. Beginning at 10 fM, a measurable increase in mean fluorescence intensity is observed for both targets, with signal levels rising progressively as the target concentration increases from 100 fM to 100 pM. Notably, Influenza H3 shows a steeper increase in fluorescence intensity compared to SARS-CoV-2 at higher concentrations, reaching approximately 80 a.u. at 100 pM, while SARS-CoV-2 reaches ~35 a.u. Fluorescence intensities for both targets display clear concentration dependence, consistent with the increased number of positive reservoirs observed qualitatively. Statistical analysis indicates strong linearity for Influenza H3 (R^2^ < 0.0001) and a weaker correlation for SARS-CoV-2 (R^2^ = 0.0009), suggesting target-dependent amplification efficiency. Overall, these results further confirm the sensitivity and quantitative capability of the duplex detection platform.

In this work, we demonstrate that the duplex RT-LAMP assay enables both colorimetric and fluorescence detection of two distinct RNA targets on a chip-scale platform. A limit of detection of 10 fM was achieved, with a dynamic range spanning five orders of magnitude. One major advantage of the chip-based LAMP system is the use of mineral oil to seal the numerous reaction reservoirs, effectively preventing cross-contamination commonly associated with conventional tube-based (e.g., Eppendorf) assays [10,11]. Furthermore, we previously demonstrated that both the LAMP reagents and mineral oil can be automatically loaded using a “vacuum lung” design, significantly simplifying operation and making the platform suitable for users without specialized laboratory training.

**Figure 4.**
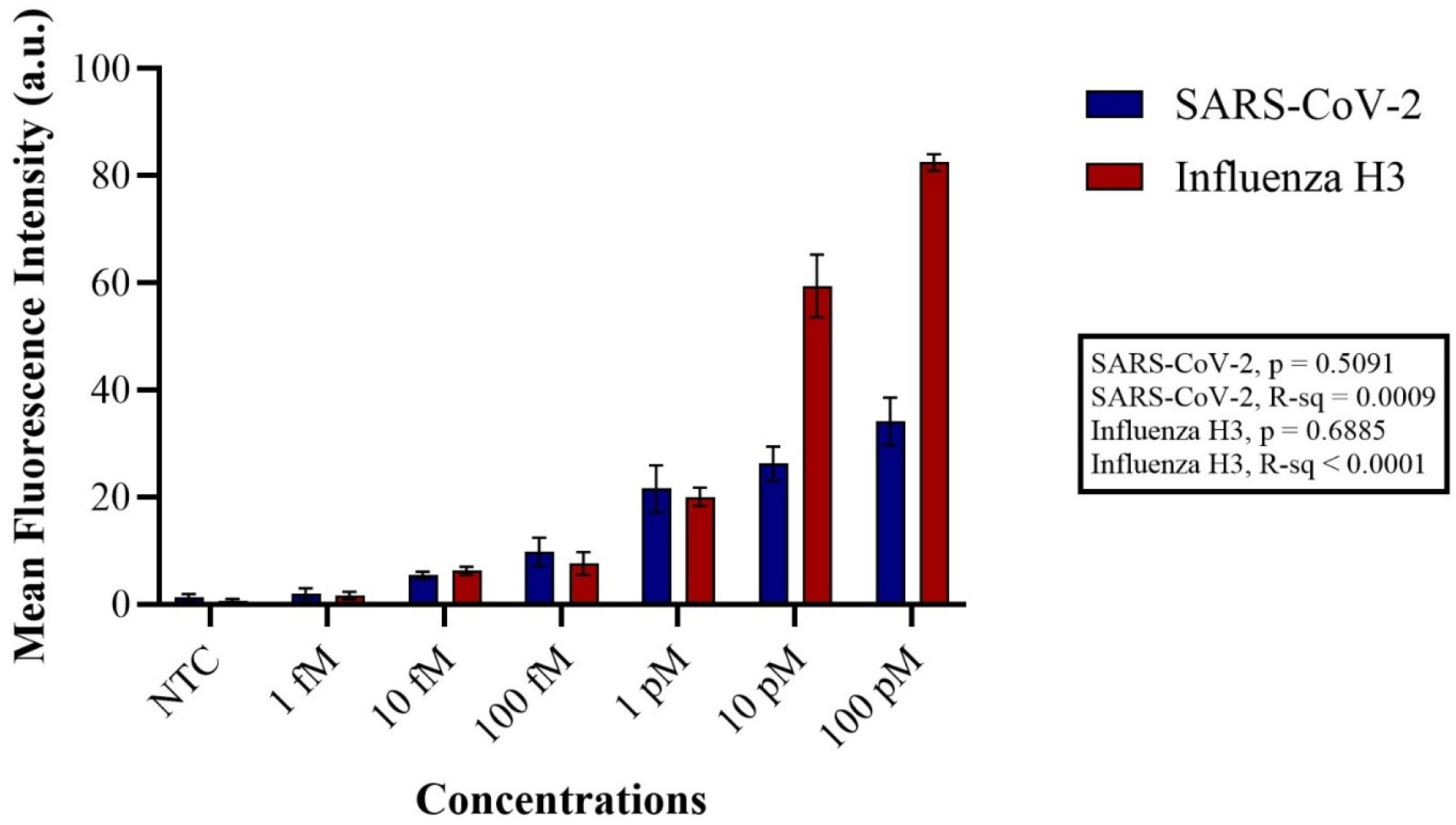
Mean fluorescence intensity of Nano-dChip reservoirs with RT-LAMP assays using SARS-CoV-2 and Influenza H3 viral RNA at six concentrations: 1 fM, 10 fM, 100 fM, 1 pM, 10 pM and 100 pM. NTC uses dH_2_O instead of target RNA.

RT-LAMP operates at a constant temperature throughout the amplification process. This isothermal approach eliminates the need for a thermal cycler, thereby significantly reducing instrument cost and operational complexity. Compared with conventional PCR, which typically requires 2–3 hours to complete, RT-LAMP can deliver results in a substantially shorter time frame—an advantage that is critical for the timely identification of pathogens [12]. We have demonstrated sensitive detection down to 10 fM (163 copies/mL) within 30 minutes. To further enhance the detection sensitivity of the Nano-dChip, the RT-LAMP assay can be integrated with CRISPR-Cas12–based signal amplification, enabling improved analytical performance and lower limits of detection [13,14].

The Nano-dChip is fabricated using standard photolithography and PDMS soft lithography techniques. Each PDMS chip costs less than $0.5, making it well suited for disposable POC diagnostics. Compared with conventional droplet microfluidic platforms, which require careful characterization and control of flow profiles at T-junctions to generate stable droplets [15], reagent loading in the Nano-dChip is significantly simpler. In addition, unlike droplet microfluidic systems that often require manual transfer of generated droplets to separate platforms for downstream quantification [16], both the RT-LAMP reaction and fluorescence readout in the Nano-dChip are performed directly on the chip, simplifying the overall workflow.

In the future, multiplexed detection of a broad panel of respiratory viruses can be achieved through further optimization of the RT-LAMP assay. For example, multicolor barcoded fluorescence labeling can replace single-color fluorophores, enabling simultaneous discrimination of multiple viruses that present with similar clinical symptoms [17]. In addition, multiplexed colorimetric detection can be implemented on the Nano-dChip by analyzing real-time signal change kinetics, allowing differentiation based on reaction rate profiles over time [18]. These enhanced assay strategies can be integrated with deep learning–based automated data analysis to improve accuracy and sensitivity [19]. The Nano-dChip is currently quantified using a commercially available fluorescence microscope. In the future, low-cost, miniaturized portable microscopy systems could be integrated with the Nano-dChip to enable field-deployable analysis [20,21]. Together, these improvements would provide reliable, high-throughput detection of multiple respiratory viral pathogens on a single compact platform while eliminating the need for bulky and complex instrumentation [22].

## Acknowledgements

This project was supported by NIH R35GM142763 and USDA NIFA 2022-67021-41478.

